# Auditory Working Memory and Sound Segregation Ability Predict Speech-in-Noise in Adult Cochlear Implant Users

**DOI:** 10.64898/2026.06.05.730315

**Authors:** Hasan Colak, Xiaoxuan Guo, Ester Benzaquén, Madhusagar Gurusiddappa, Anirvan Banerjee, Inyong Choi, William Sedley, Timothy D. Griffiths

**Affiliations:** Biosciences Institute, Newcastle University, Newcastle upon Tyne, NE2 4HH, United Kingdom; Department of Audiology, Hacettepe University, Ankara, Türkiye; Instituto de Neurociencias de Alicante, CSIC-UMH, San Juan de Alicante, Spain; James Cook University Hospital, South Tees Hospitals NHS Foundation Trust, Middlesbrough, United Kingdom; Department of Communication Sciences and Disorders, University of Iowa, Iowa City, IA, USA; Translational and Clinical Research Institute, Faculty of Medical Sciences, Newcastle University.; Department of Imaging Neuroscience, University College London, London, UK

**Keywords:** Cochlear implant, speech-in-noise, auditory working memory, auditory figure ground, sound segregation, auditory processing, speech perception

## Abstract

**Objectives:** Outcomes following cochlear implantation vary substantially across adult recipients, and the cognitive and perceptual factors contributing to this variability are not fully understood. This poses a challenge for developing strategies to improve cochlear implant outcomes, as such approaches require a clearer understanding of the mechanisms underlying individual listening difficulties. In this study, we investigated auditory cognitive measures in cochlear implant (CI) users to further elucidate the origins of this variability.

**Design:** Thirty-seven adult cochlear implant users completed measures of auditory cognition, comprising auditory working memory (AWM) and sound segregation ability, measured using an auditory figure–ground task (AFG), as well as measures of peripheral temporal and spectral processing, comprising the temporal modulation detection threshold (TMDT) and spectral ripple discrimination threshold (SRDT). Speech perception outcomes were assessed using word-in-noise (WIN) and sentence-in-noise (SIN) tasks. Separate multiple linear regression models evaluated the unique contribution of the auditory cognition measures to WIN and SIN performance, after accounting for the peripheral measures.

**Results:** Both regression models explained a substantial proportion of variance in speech-in-noise outcomes (WIN: adjusted R² = 0.55; SIN: adjusted R²=0.57, both *p* < 0.001). For WIN performance, AFG and AWM were significant predictors. A similar pattern was found for SIN performance, where lower AWM ability and poorer AFG segregation were linked to poorer sentence listening in noise. No significant effects of spectral ripple discrimination or temporal modulation detection were observed in either model, even though both were significantly correlated with WIN performance.

**Conclusions:** These findings indicate that auditory working memory and sound segregation ability are robust predictors of speech-in-noise outcomes in adult cochlear implant users, across both word- and sentence-level measures. Together, the results may help explain why speech-in-noise outcomes remain highly variable among CI users, even when basic sensory encoding abilities are taken into account. Incorporating measures of auditory working memory and fundamental sound segregation may therefore improve outcome prediction and help in developing more individualised rehabilitation strategies.

## Introduction

To date, cochlear implants (CIs) have been one of the most successful neural prostheses that help people with severe-to-profound hearing loss to gain or regain the ability to hear, making verbal daily life communication possible. Although the devices perform well in speech understanding in quiet environments, speech perception in noise remains a significant challenge for CI users. Importantly, speech-in-noise performance varies considerably across CI users, even among those with similar audiological profiles, suggesting that factors beyond peripheral hearing restoration contribute to this variability (Holden et al., 2013). This is consistent with large-sample studies showing that demographic and audiological factors explain only a limited proportion of variance in speech perception outcomes, even for speech perception in quiet (Blamey et al., 2013; Goudey et al., 2021). Given that most daily life activities involve background noise, whether from multiple simultaneous talkers or competing environmental sounds, it is crucial to understand the mechanisms of speech-in-noise in CI users so that effective rehabilitation strategies and device optimisation approaches can be developed.

Speech-in-noise difficulty can even be seen in people with normal pure tone hearing thresholds (Smith et al., 2019), suggesting that peripheral hearing does not fully account for speech-in-noise ability. Real-world listening requires listeners to detect and follow relevant speech patterns within a complex and changing auditory background, indicating higher-level auditory processing involvement rather than a problem of audibility alone. From first principles, speech-in-noise ability requires auditory stream formation and successful sound segregation through the grouping and temporal integration of acoustic elements over time. Auditory figure–ground (AFG) perception captures this ability by measuring how well listeners detect coherent tonal elements embedded within randomly varying background tones (Teki et al., 2013). Previous work on AFG suggested that this high-level, fundamental sound-grouping ability is a significant predictor of speech-in-noise performance in normal hearing listeners and explains variance in speech-in-noise performance independent of that explained by hearing thresholds (Benzaquén et al., 2025; Guo et al., 2025; Holmes & Griffiths, 2019; Holmes et al., 2021). A recent study by Choi et al. (2023) demonstrated a significant contribution of AFG ability to speech-in-noise performance, mostly in hybrid CI users in which both electrical and acoustic hearing are provided within the same ear, which was independent of peripheral spectral and temporal resolution. The model they proposed, including spectral and temporal resolution measures, significantly predicted 46% of the variance in a sentence-in-noise listening task.

Successful speech-in-noise perception depends not only on segregating relevant auditory information from the background, but also on maintaining and updating that information over time. Once speech cues are grouped into a coherent auditory stream, listeners must hold partial or degraded acoustic information in memory while integrating it with subsequent input. This temporary maintenance and recall of relevant auditory information place demand on auditory working memory (AWM). The effect of working memory on speech-in-noise perception has been the subject of extensive research in normal-hearing listeners. Although some studies have reported weaker or null effects (e.g., Füllgrabe and Rosen (2016)), the majority have suggested that better working memory abilities are associated with better speech-in-noise performance (Akeroyd, 2008; Benzaquén et al., 2025; Kraus, Strait, & Parbery-Clark, 2012; Lad et al., 2020; Rönnberg et al., 2013; Yeend, Beach, & Sharma, 2019). The relationship between AWM and speech-in-noise performance has also been investigated in CI users, but the findings have mostly indicated no significant association between AWM and speech-in-noise abilities (Beckers et al., 2023; Luo et al., 2022; Moberly, Harris, et al., 2017; Tao et al., 2014). Nonetheless Moberly, Houston, et al. (2017) reported that performance on a listening span task, but not a reading span task, was significantly related to speech recognition in adult CI users. However, it is important to note that previous studies in CI users mostly suffered from small sample sizes and used verbal auditory working memory tasks, such as digit- or word-based measures. Performance on such tasks inherently depends on further linguistic processes such as phonological encoding and lexical access, which are themselves compromised by the spectrally degraded input provided by CIs. Despite this lack of convincing evidence, AWM may in fact play an even greater role for CI users than for normal-hearing listeners, given that they experience substantially greater difficulty understanding speech in adverse listening conditions and may therefore place greater reliance on cognitive resources such as working memory to compensate for degraded auditory input. It is possible, then, that these null findings reflect the confounded nature of the tasks used rather than a genuine absence of an AWM contribution in CI users. To address this, we adopted an AWM task that relies mainly on temporal cues such as amplitude modulation (AM) rather than spectral or linguistic information, since AM encoding is relatively well preserved in CI users compared with spectral encoding (Zeng, 2022; Zhou et al., 2020).

The aim of the present study was to investigate the auditory cognitive contributions to speech-in-noise perception in adult CI users with standard long-electrode arrays covering both the basal and apical regions of the cochlea, thereby avoiding the confound of residual acoustic hearing present in hybrid CI users. Specifically, we examined the contributions of two higher-level auditory abilities, AFG perception and AM-based AWM, to speech-in-noise performance. These were assessed alongside measures dependent on spectral and temporal encoding, which together provide a complementary account of sensory encoding quality at peripheral level. We investigated whether AFG perception and AM-based auditory working memory explain variance in speech-in-noise performance over and above that accounted for by spectral and temporal resolution.

## Materials and Methods

### Participants

This study initially included 40 adult cochlear implant users recruited from the North East Regional Cochlear Implant Programme in the United Kingdom. All participants had postlingual severe-to-profound hearing loss prior to cochlear implantation, were native English speakers, and had no neurological or psychological comorbidities. All participants fulfilled the pre-implantation NICE criteria for cochlear implantation, which define severe-to-profound bilateral deafness as unaided hearing thresholds of ≥80 dB HL at two or more frequencies across 0.5, 1, 2, 3, and 4 kHz. All participants were conventional cochlear implant users, meaning that they received only electric stimulation rather than a combination of acoustic and electric stimulation within the same ear, as in hybrid cochlear implants. To be included in the study, participants had to be over 18 years of age, use a unilateral cochlear implant, and have at least one year of cochlear implant experience. Exclusion criteria included bilateral cochlear implantation, prelingual hearing loss, incomplete data, and neurological or psychological comorbidities. One participant with bilateral cochlear implants was excluded, and two additional participants were excluded because of incomplete data. The final sample therefore consisted of 37 unilateral cochlear implant users (18 female), aged 22 to 81 years, with a mean age of 60.62 years (SD = 17.07). Seventeen participants used a left CI and 20 used a right CI. Most participants used Advanced Bionics devices (n = 27), followed by Cochlear/Nucleus devices (n = 8). One participant used a MED-EL device and one used an Oticon device. Some participants also used a hearing aid in the non-implanted ear, but they were asked to remove it or switch it off during testing to ensure that acoustic input from that ear did not affect the results. All testing took place in a soundproof room and was performed by a trained audiologist. Custom computerized auditory tests were administered using two loudspeakers (Yamaha HS8 Active Studio Monitors) positioned at ±45° azimuth relative to the listener. Stimuli were presented using Psychtoolbox in MATLAB 2017a. Practice trials were given for all tasks to familiarize participants with the task and stimulus. All tasks were conducted using the listeners’ common listening mode for the device used in their daily life. This study received a favourable opinion from the London - Central Research Ethics Committee (IRAS ID: 311586; REC reference: 22/PR/1684), and written informed consent was obtained from all participants before participation.

### Speech-in-Noise Listening

Speech-in-noise testing was conducted at both the single-word and sentence levels. The British version of the Iowa Test of Consonant Perception was used to assess single-word recognition in noise (Guo et al., 2024). The test consists of phonetically balanced single words presented against 8-talker babble. Words were spoken by one female and one male speaker, both using a standard Southern British accent. The task consisted of 120 trials, in which the target words were monosyllabic and followed a consonant–vowel–consonant (CVC) structure. Participants heard a target word presented in background noise and were shown four word options on the screen, from which they had to select the target word by pressing the corresponding number on the keyboard (e.g., “shore–more–pour–tore”). All target words were presented at a fixed signal-to-noise ratio (SNR) of +15 dB for all participants, and the speech level was set at 70 dB SPL. Performance was quantified as the percentage of correctly identified target words out of 120 trials.

Sentence-level speech-in-noise ability was assessed using the English version of the Oldenburg sentence test. The target sentences followed a fixed five-word structure: <NAME> <VERB> <NUMBER> <ADJECTIVE> <NOUN> (e.g., Doris gives fifteen heavy chairs). Sentences were presented against a 16-talker babble background, as in the original study (Holmes & Griffiths, 2019). Unlike the word-in-noise task, this task followed an adaptive 1-down, 1-up procedure. The starting SNR was +20 dB, and this level was adaptively adjusted based on participant’s performance. The target speech level remained constant at 70 dB SPL, while the noise level was varied according to performance. A response was considered correct only when all five words were identified correctly. Participants were shown a 5 × 10 matrix and were required to select the words they heard to build the sentence by clicking on the appropriate choices in each word category. The total number of reversals was 10, and the step size began at 2 dB and decreased to 0.5 dB after three reversals. The task comprised two interleaved runs, and different target sentences were presented in each run. The final score was calculated by averaging the SNR values at the last six reversals across the two runs. Lower scores indicated better performance on this test.

### Auditory Figure-Ground (AFG) Detection

Participants completed an AFG detection task, adapted from the paradigm introduced by Teki et al. (2013), to assess their ability to segregate sound patterns from a random acoustic background. Each tone chord lasted 50 ms, and the possible frequencies used to generate the sound sequences ranged from 180 to 7246 Hz. Some of these tone chords repeated at fixed frequencies over time, creating the ‘figure’, whereas others remained random and formed the background component. During the task, participants heard either a figure sound embedded in random background tone clouds or a ground-only sound consisting solely of random background tone clouds. The coherence level was set to 6, meaning that six different frequencies remained constant over time in the figure sounds. The number of background components was fixed at 8. In each trial, participants heard a 4-second sound sequence. In ground-only trials, the entire sequence consisted of random tone clouds. In figure trials, the first 2 seconds consisted only of the ground component, whereas during the final 2 seconds both figure and ground components were presented together. An illustration of both figure and ground sounds is provided in Figure 1. Additional normalisation was applied to the figure-embedded sound complexes to prevent the final part of the sequence from having higher overall energy because of the combination of figure and ground components. After each sound, participants were asked to indicate whether they heard a target sound using a simple yes/no keyboard response. The task included 120 trials, half of which contained a figure and half of which consisted of ground-only sounds. Before the main task, 18 practice trials were presented, and example spectrograms of figure and ground sounds were shown visually to familiarise participants with the task. Performance was quantified using d′. The sound intensity for both figure and ground trials was kept at 70 dB SPL.

**Figure 1.**
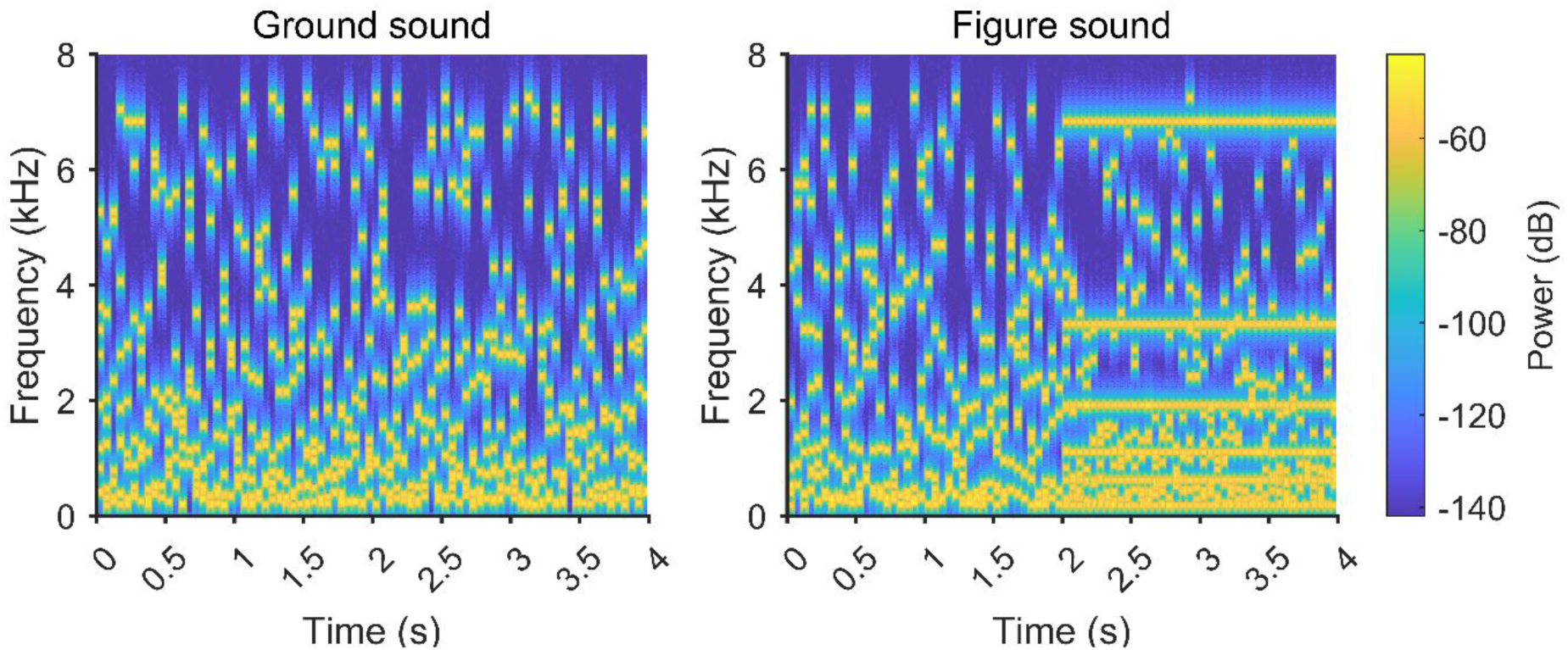
Spectrograms of example auditory figure–ground stimuli. Spectrograms showing the time–frequency structure of representative ground and figure sounds. The ground sound consists of randomly varying tonal components distributed over time and frequency. The figure sound contains the same type of background structure but includes a temporally coherent figure pattern, visible as repeated horizontal frequency components emerging from the background in the second half of the stimulus. Frequency is shown in kHz, time in seconds, and colour indicates sound power in dB.

### Auditory Working Memory (AWM)

Auditory working memory was assessed using a task that required participants to hold amplitude-modulated white noise in memory. In our previous work, we used a similar task with both tones and amplitude-modulated white noise in normal-hearing listeners (Lad et al., 2022; Lad et al., 2020). Given that frequency encoding is substantially impaired in CI users, whereas temporal processing abilities are relatively preserved, only amplitude-modulated sounds were used in the present study. In the task, participants first heard a target sound consisting of amplitude-modulated white noise and were asked to remember it. After a 2-s silent delay, a horizontal slider appeared on the screen. Participants were instructed to recreate the remembered AM rate as accurately as possible by moving the slider with the mouse. This procedure required participants to keep the target sound in memory while comparing it with newly generated sounds, thereby engaging auditory working memory. Both the target sound and the sounds generated during the matching phase were sinusoidally amplitude-modulated white noise with 100% modulation depth and a duration of 0.75 seconds. The modulation rate of the target was randomly selected from a range of 5–30 Hz. The starting phase of the AM modulator was held constant across trials. The slider was mapped onto a 20-Hz range, which was randomly positioned on each trial while always including the target modulation rate, in order to reduce learning of fixed slider positions. Each slider movement triggered a new 0.75-s amplitude-modulated noise stimulus corresponding to the selected slider position, allowing participants to compare their current choice with the remembered target. Participants were informed only that lower AM rates would be presented on the left side of the slider and higher AM rates on the right side. After the slider appeared, participants were given 15 seconds to identify the target sound and complete their adjustment. After each trial, participants received brief feedback to help maintain engagement with the task. This feedback indicated the percentage distance between the participant’s selected modulation rate and the target modulation rate. Lower percentage values reflected a closer match. When the selected value was within 2% of the target, participants were shown the message “Exact target match!”. This feedback was intended only to encourage engagement and was not used as an outcome measure. Participants completed 120 trials in the main task, and 12 separate practice trials were presented beforehand. For each trial, matching error was measured as the difference between the target value and the participant’s selected value. Precision was calculated as the inverse of the standard deviation of the matching errors across trails. The precision values were log transformed to allow parametric statistics which were the main outcome of the task. The sounds were presented at 70 dB SPL using loudspeakers. An example sound and the trial structure of the task are shown in Figure 2.

**Figure 2.**
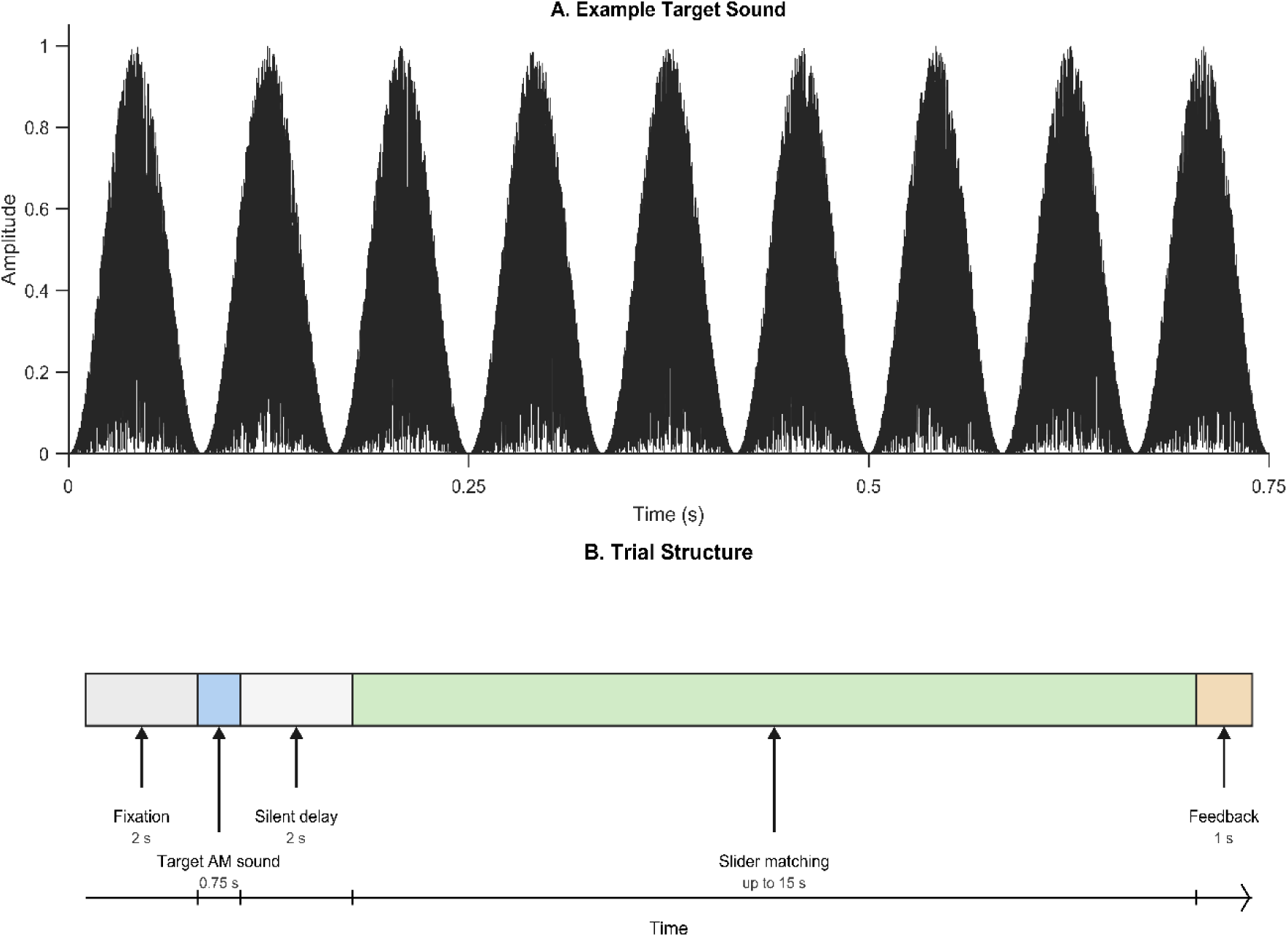
Auditory working memory task. A. Example target sound used in the task. The stimulus was a 0.75-s amplitude-modulated noise sound at 12 Hz. Participants were required to remember this modulation rate during the trial. B. Schematic of the trial structure. Each trial began with a 2-s fixation period, followed by presentation of the 0.75-s target sound. After a 2-s silent delay, participants adjusted a slider to reproduce the remembered modulation rate. Each slider movement triggered a new 0.75-s amplitude-modulated sound corresponding to the selected slider position. The response period lasted up to 15 s and was followed by 1 s of feedback.

### Spectral Ripple and Temporal Modulation Tasks

Spectral ripple discrimination and temporal modulation detection thresholds were measured using a three-interval oddball discrimination task. On each trial, participants heard three sounds, two of which were standard stimuli and one of which was the target stimulus. The target interval was randomly assigned on each trial. Participants were asked to indicate which of the three intervals contained the sound that was different by pressing the corresponding number key or selecting the corresponding response box on the screen. Visual feedback was provided after each response. Stimuli were generated in MATLAB and presented using Psychtoolbox. Each stimulus had a duration of 500 ms, and the inter-stimulus interval was 750 ms. The root mean square level of each stimulus was normalised prior to presentation, and an independent ±3 dB level rove was applied to each interval to minimise the use of overall level cues. Thresholds were estimated using an adaptive procedure implemented with the Updated Maximum-Likelihood (UML) adaptive track engine. Following 4 practice trials, participants completed 70 experimental trials for each condition. The adaptive procedure updated the stimulus value after each response and estimated the psychometric parameters across trials. The final threshold was taken from the estimated crossover value at the end of the adaptive track. In the spectral ripple task, the stimuli were broadband noise signals shaped by sinusoidal modulation along a logarithmic frequency axis. The ripple density was kept constant at 1.25 ripples per octave. Ripple depth, measured in dB, was adjusted adaptively using the UML procedure. Ripple depth, measured in dB, was adjusted adaptively using the UML procedure.

For each trial, the two standard intervals were generated with a randomly selected spectral ripple phase. The task measured participant’s spectral ripple depth thresholds by adaptively varying ripple depth to estimate the minimum depth required to identify the oddball interval. For the temporal modulation detection task (TMDT), stimuli were generated using seven carrier frequencies: 332, 551, 913, 1517, 2519, 4183, and 6937 Hz. The envelope modulation frequency was fixed at 20 Hz. Modulation depth was varied adaptively to estimate each participant’s temporal modulation detection threshold. On each trial, participants heard either two modulated stimuli and one unmodulated oddball, or two unmodulated stimuli and one modulated oddball. The participant’s task was to identify the interval that differed in temporal modulation from the other two intervals.

### Statistical Analysis

Statistical analyses were performed in MATLAB R2024b. Descriptive statistics were first calculated for all behavioural and auditory-cognitive measures. Spearman’s correlation coefficients were first calculated to examine bivariate associations between speech-in-noise performance and potential predictor variables, including auditory figure–ground performance, auditory working memory. spectral ripple depth thresholds, and temporal modulation detection thresholds. To further examine the contribution of auditory and cognitive predictors to speech-in-noise perception, multiple linear regression models were fitted separately for word- and sentence-level speech-in-noise performance using the fitlm function in MATLAB. The word-in-noise (WIN) model included spectral ripple depth threshold, temporal modulation detection threshold, auditory figure–ground performance, and auditory working memory as predictors. The same predictor structure was used for the sentence-in-noise (SIN) model. The model formulae were:

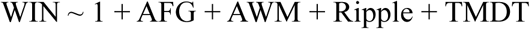

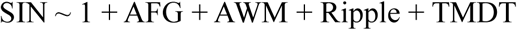

Prior to model fitting, variance inflation factors were calculated to assess multicollinearity among predictors. All VIF values were ≤ 1.53, suggesting that multicollinearity was not a concern for this dataset. Statistical significance was determined using an alpha level of 0.05.

## Results

Descriptive statistics and participant characteristics are presented in Table 1.

**Table 1.**
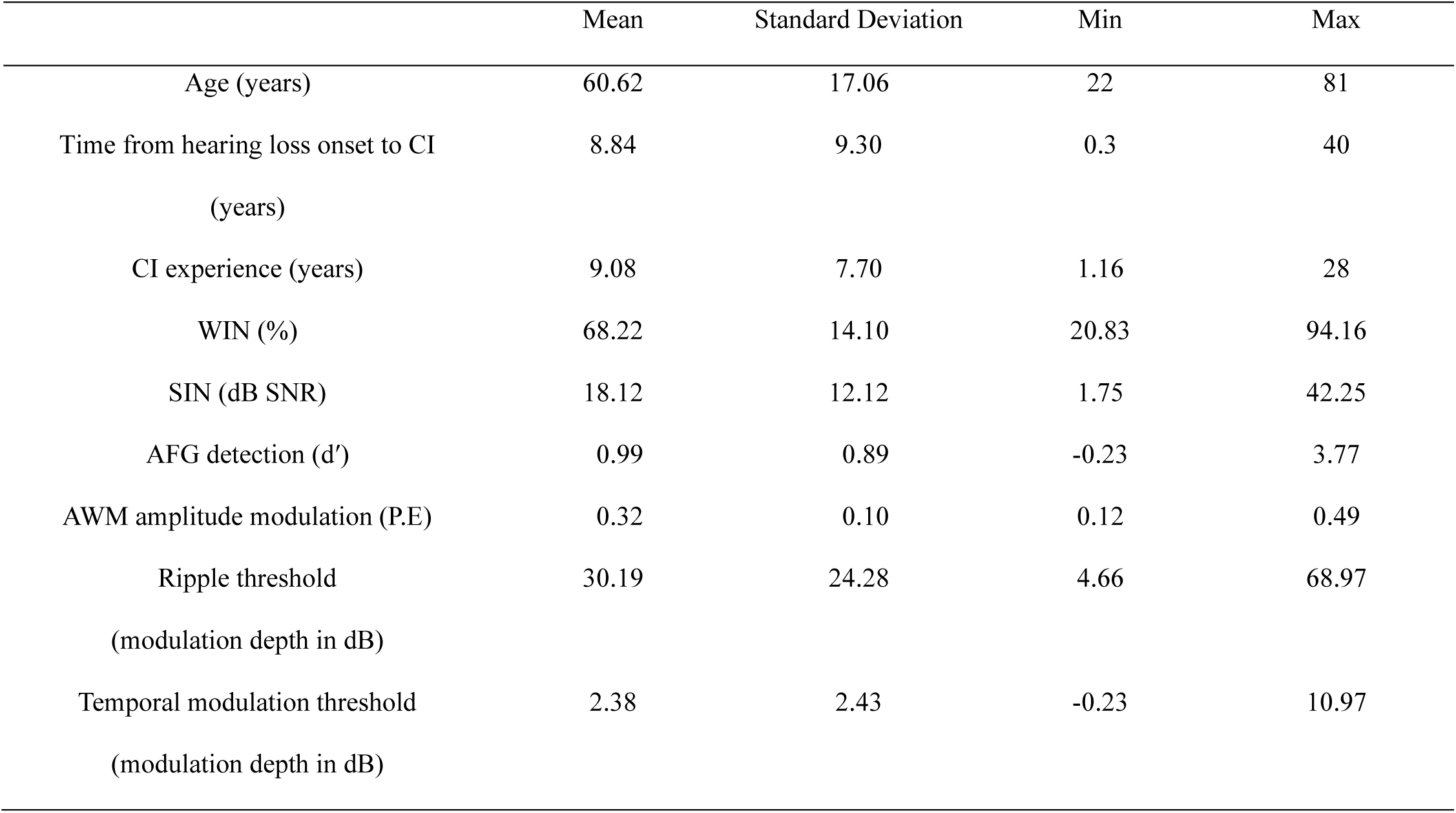
Participant demographics and task performance. Values represent mean, standard deviation, minimum, and maximum. WIN = word-in-noise; SIN = sentence-in-noise; AFG = auditory figure–ground; AWM = auditory working memory; P.E. = precision estimate. Higher WIN, AFG, and AWM values indicate better performance, whereas lower SIN, ripple threshold, and temporal modulation threshold values indicate better performance.

### Correlates of Speech-in-Noise Listening

Spearman’s correlations were first used to examine bivariate associations between speech-in-noise performance and auditory-cognitive measures (Figure 3). WIN performance was significantly associated with both AFG detection and AWM. Better WIN performance was associated with higher AFG sensitivity, ρ = 0.77, p < 0.001, and higher AWM precision, ρ = 0.51, p = 0.001. These associations corresponded to large and moderate-to-large effects, respectively. SIN performance showed a similar pattern, although in the opposite direction because lower SIN thresholds indicate better performance. Better SIN performance was associated with higher AFG sensitivity, ρ = −0.68, p < 0.001, and higher AWM precision, ρ = - 0.63, p < 0.001, both corresponding to large effects.

**Figure 3.**
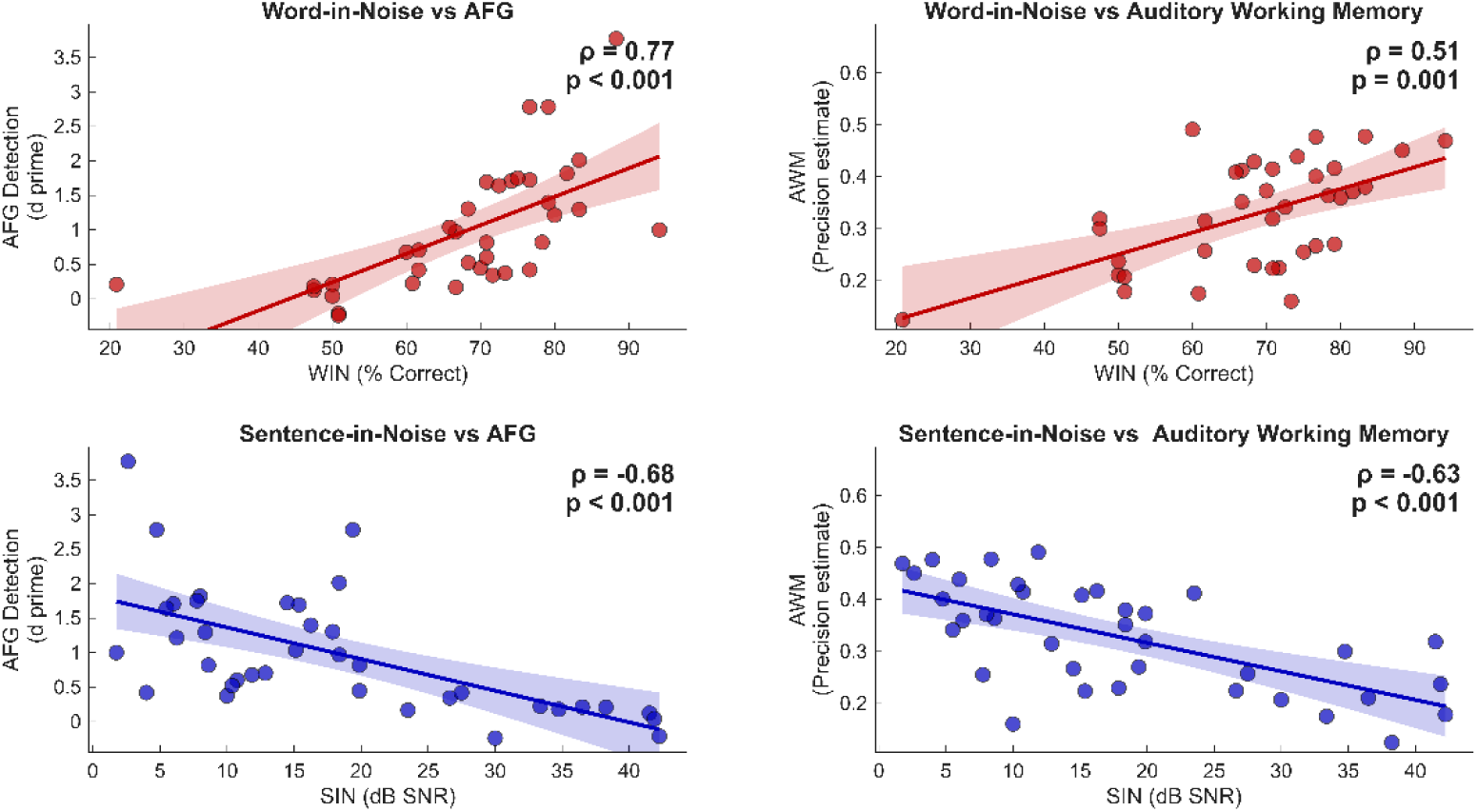
Associations between speech-in-noise performance and auditory-cognitive measures. Scatterplots show relationships between word-in-noise performance (WIN; % correct) and auditory figure–ground detection (AFG; d′) and auditory working memory (AWM; precision estimate), shown in red in the top row. The bottom row shows associations between sentence-in-noise performance (SIN; dB SNR) and the same auditory-cognitive measures, shown in blue. Lines represent linear fits for visualisation, with shaded areas indicating 95% confidence intervals. Spearman’s correlation coefficients (ρ) and associated p-values are displayed in each panel. Higher WIN scores indicate better word recognition, whereas lower SIN thresholds indicate better sentence-in-noise performance.

We also examined associations between speech-in-noise performance and spectral and temporal resolution measures (Figure 4). WIN performance was negatively correlated with both spectral ripple threshold, ρ = −0.49, p = 0.002, and temporal modulation threshold, ρ = - 0.49, p = 0.002. These moderate effects indicate that better WIN performance was related to lower, and therefore better, spectral ripple and temporal modulation thresholds. For SIN performance, correlations with spectral ripple threshold, ρ = 0.30, p = 0.067, and temporal modulation detection threshold, ρ = 0.31, p = 0.064, showed positive trends but did not reach statistical significance.

**Figure 4.**
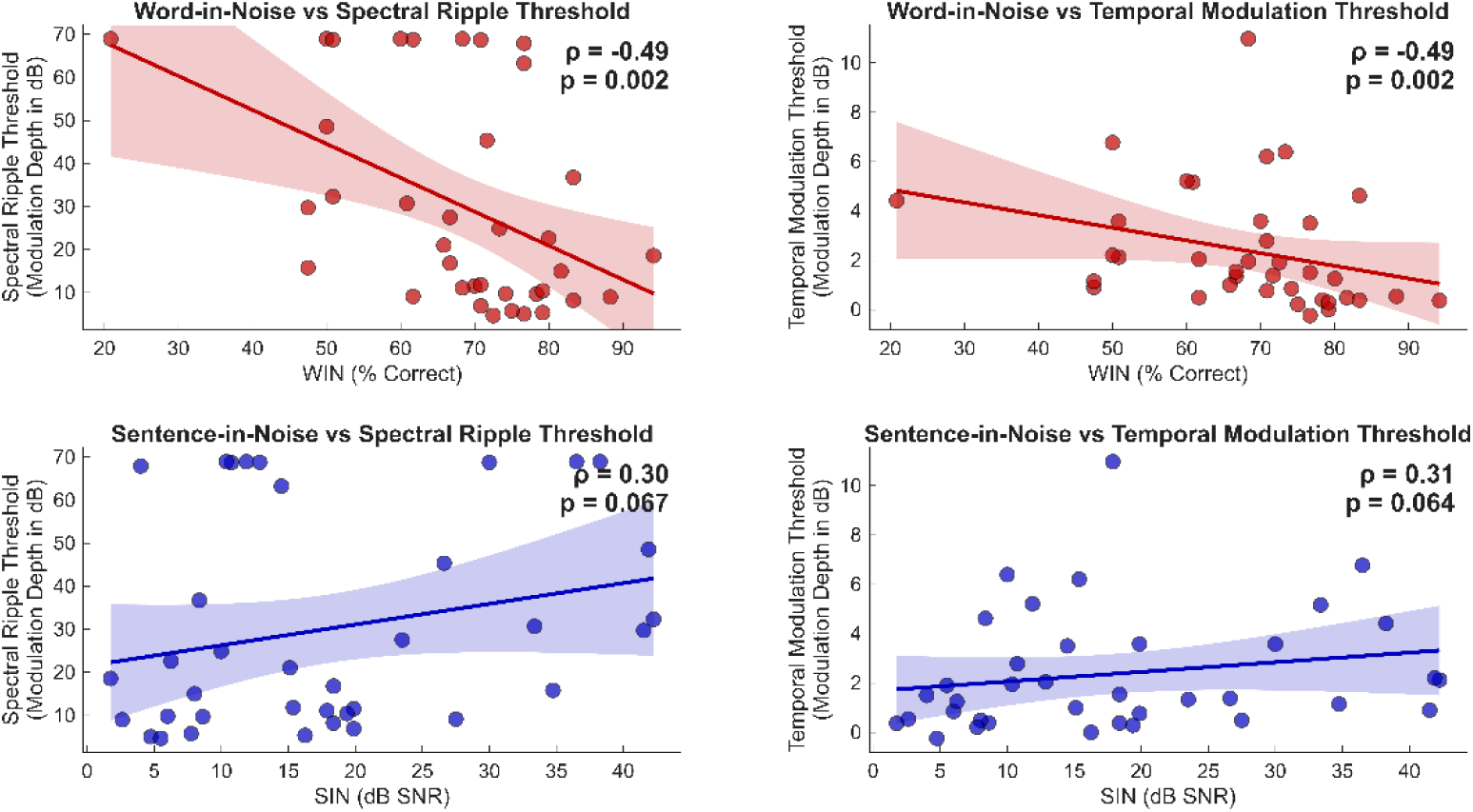
Relationship between speech-in-noise performance and spectral/temporal resolution. Word-in-noise scores (WIN; % correct) are plotted against spectral ripple and temporal modulation thresholds in the upper panels, while sentence-in-noise thresholds (SIN; dB SNR) are plotted against the same measures in the lower panels. Red markers indicate WIN associations and blue markers indicate SIN associations. Solid lines show fitted linear trends, and shaded regions represent 95% confidence intervals. Spearman’s correlation coefficients (ρ) and corresponding p-values are shown within each panel. Lower spectral ripple and temporal modulation thresholds reflect better spectral and temporal resolution, respectively; higher WIN scores and lower SIN thresholds reflect better speech-in-noise performance.

### Predictors of Speech-in-Noise Listening

Multiple linear regression models were then fitted separately for WIN and SIN performance to estimate the combined contribution of these potential predictors of speech-in-noise ability. For WIN performance, the overall model was statistically significant, F(4, 32) = 12.10, p < 0.001, and explained 60.3% of the variance in word-in-noise scores, R² = 0.603, adjusted R² = 0.553. AWM was a significant positive predictor of WIN performance, β = 0.622, SE = 0.180, t = 3.46, p = 0.002. AFG detection was also a significant predictor, β = 0.062, SE = 0.022, t = 2.84, p = 0.008. In contrast, temporal modulation detection threshold was not a significant predictor, β = 0.004, SE = 0.007, t = 0.53, p = 0.602. Spectral ripple threshold showed a trend but did not reach statistical significance, β = −0.001, SE = 0.001, t = −1.74, p = 0.092.

For SIN performance, the overall model was also statistically significant, F(4, 32) = 13.30, p < 0.001, and explained 62.4% of the variance in sentence-in-noise thresholds, R² = 0.624, adjusted R² = 0.577. AWM was a significant predictor of SIN thresholds, β = −66.39, SE = 15.04, t = −4.41, p < 0.001. AFG detection was also a significant predictor, β = −6.35, SE = 1.81, t = −3.50, p = 0.001, indicating that better AFG performance was associated with lower, and therefore better, SIN thresholds. Temporal modulation detection threshold did not significantly predict SIN performance, β = −0.81, SE = 0.63, t = −1.29, p = 0.206, and spectral ripple threshold was also not a significant predictor, β = −0.002, SE = 0.066, t = −0.02, p = 0.982. The regression models for both WIN and SIN performance are presented in Table 2.

**Table 2.** Multiple linear regression models predicting word- and sentence-in-noise performance. WIN = word-in-noise; SIN = sentence-in-noise; AFG = auditory figure–ground; AWM = auditory working memory; TMDT = temporal modulation detection threshold; RMSE = root mean squared error. β values are unstandardised regression coefficients.

| Speech-in-noise Outcome | Predictor | $\beta$ | SE | t | p |
| --- | --- | --- | --- | --- | --- |
| WIN | Intercept | 0.449 | 0.069 | 6.51 | < .001 |
|  | AFG detection | 0.062 | 0.022 | 2.84 | .008 |
|  | AWM | 0.622 | 0.180 | 3.46 | .002 |
|  | Ripple threshold | -0.001 | 0.001 | -1.74 | .092 |
|  | Temporal modulation threshold | 0.004 | 0.007 | 0.53 | .602 |
| SIN | Intercept | 48.127 | 5.772 | 8.34 | < .001 |
|  | AFG detection | -6.347 | 1.814 | -3.50 | .001 |
|  | AWM | -66.389 | 15.039 | -4.41 | < .001 |
|  | Ripple threshold | -0.002 | 0.066 | -0.02 | .982 |
|  | Temporal modulation threshold | -0.806 | 0.625 | -1.29 | .206 |
| Model Fit | RMSE | R<sup>2</sup> | Adjusted R<sup>2</sup> | F | p |
| WIN | 0.094 | 0.603 | 0.553 | F(4, 32) = 12.10 | < .001 |
| SIN | 7.89 | 0.624 | 0.577 | F(4, 32) = 13.30 | < .001 |

Overall, the regression analyses suggested that AWM and AFG detection were the strongest independent predictors of both word- and sentence-level speech-in-noise performance. These effects remained significant even after accounting for spectral and temporal resolution, which are considered highly relevant performance indicators in CI listeners.

## Discussion

In this study, we investigated whether higher-level auditory processing abilities, particularly fundamental sound-grouping ability measured with the AFG task and the ability to hold sounds in mind measured with the AWM task, can predict speech-in-noise outcomes in conventional CI users. We observed substantial variability in speech-in-noise performance, as well as in other measures of basic sensory processing and higher-level auditory processing. In terms of bivariate correlations, both AFG and AWM performance showed significant correlations with word- and sentence-level speech-in-noise tests, with large effect sizes. Further, linear regression models demonstrated that AFG and AWM abilities were significant predictors of both word- and sentence-level speech-in-noise listening, even after accounting for peripheral spectral and temporal resolution performance. Models including AFG and AWM, together with these fundamental sensory processing indices, explained 55.3% and 57.7% of the variance in word-in-noise and sentence-in-noise performance, respectively.

The significant contribution of the ability to segregate sounds and group them into coherent elements highlights the importance of successful auditory object formation in CI users. This ability is likely to depend on the integration of spectral and temporal information, allowing the listener to form a coherent auditory object, referred to here as the “figure.” Therefore, it was important to control for CI sensory encoding fidelity using spectral and temporal resolution tests. In everyday listening, processes similar to those engaged during AFG may be required to segregate target speech from competing background sounds. Therefore, successful AFG segregation may capture a more fundamental, language-independent auditory organization process in the brain that supports speech-in-noise listening (Holmes et al., 2021). The significant contribution of AFG, together with the language-independent nature of the task, may hold a first step towards its clinical use. For example, AFG performance could be used to track and monitor patient progress over time. In addition, the AFG stimuli could be adjusted so that the figure elements occur within specific frequency ranges, or across two different devices, such as a CI and a hearing aid, allowing perceptual fusion across devices to be tested (Choi et al., 2023). The contribution of AFG to speech-in-noise listening in the present study appears to be larger than previously reported by Choi et al. (2023). One possible explanation for this larger effect size is that the current sample included only users of conventional CIs with long electrodes, unlike the sample in Choi et al. (2023). This may have reduced the confounding influence of residual acoustic hearing in the present study. Another possible explanation is that our AFG task included a more complex background tone cloud, with a greater number of background elements than in the earlier study. This may have created a more realistic listening experience, where multiple competing tonal elements are present in the background. As a result, the task may have been more representative of real-world listening demands.

We also demonstrated a significant effect of AWM for AM rate on both word- and sentence-level speech-in-noise ability. Intuitively, working memory may be especially relevant for sentence-level listening, where individual word units need to be temporarily maintained, grouped, and linked together. This process requires phonological storage, as well as a processing component that allows relevant information to be accessed and used when needed (Baddeley, 2012; Conway et al., 2005). This interpretation is supported by the larger effect sizes observed for sentence-in-noise performance in both the correlation analyses and the linear regression models. However, listening to words in noise may still involve cognitive processes such as sustaining selective attention and inhibitory control, which could explain the significant correlation between AWM and word-in-noise performance, as well as the significant contribution of AWM to word-in-noise listening in the regression models. The present study therefore addresses an important gap in the existing literature by demonstrating the relevance of AWM, measured using non-verbal stimuli, for speech-in-noise listening in CI users.

### Clinical Implications and Future Work

The findings from this study may help explain why some CI users perform well in quiet conditions but continue to struggle in noisy environments. A person may have adequate audibility through their CI and relatively good speech perception scores in quiet, but still have poor auditory grouping or AWM ability, which may limit communication in real-world settings. These results may therefore have important implications for cochlear implant fitting and rehabilitation strategies. For example, the language-independent nature of the AFG task may allow clinicians to compare results across different populations more easily. It may also help bypass some complex linguistic processes that can be affected by secondary factors, such as ageing. This suggests that the AFG task could potentially be used in clinical settings as a complement to standard speech perception test batteries. Alternatively, the AFG concept could be adapted into a training tool aimed at improving speech-in-noise listening in everyday environments. In such a training approach, listeners could practise detecting and tracking a target figure embedded within background elements, with the level of difficulty adjusted progressively. By repeatedly challenging the listener’s ability to segregate sounds and form coherent auditory objects, this type of training may then translate into better speech-in-noise performance. Our future work will focus on developing effective training strategies based on the AFG concept, first in normal-hearing listeners and then in clinical populations where speech-in-noise difficulties are common, such as CI and hearing aid users. A similar training approach could also target higher-level auditory cognition, with AWM as the main focus. The aim would be to strengthen auditory cognitive resilience and help compensate for the reduced sound quality provided by the CI.

### Limitations

There are several limitations that should be considered. First, although a sample of 37 CI users is not small for a CI study, it may still be limited for multivariate analyses based on four predictive variables. This could lead to either overestimation or underestimation of the linear regression results. It may also make it more difficult to detect smaller independent effects of spectral or temporal resolution, both of which have previously been shown to be relevant to speech-in-noise ability (Aronoff et al., 2021; Blankenship, Meinzen-Derr, & Zhang, 2022; Gifford et al., 2018; Harris et al., 2023; Lawler, Yu, & Aronoff, 2017; Won, Drennan, & Rubinstein, 2007). Second, although we attempted to control for peripheral sensory encoding by including spectral and temporal resolution measures, the AFG and AWM tasks may not fully overlap with these measures in terms of stimulus characteristics or the specific peripheral processes they engage. Therefore, there may still be some contribution of basic sound encoding to performance on the AFG and AWM tasks in CI users.

## Conclusion

In summary, our results demonstrate significant contributions of both AFG and AWM abilities to word- and sentence-level speech-in-noise performance in adults with post-lingual hearing loss who use CIs. These contributions were independent of spectral and temporal resolution abilities, which were included as measures of sensory processing with the CI. These findings suggest that difficulties in segregating sounds and grouping them into coherent auditory objects, together with poor AWM ability, may be among the mechanisms underlying poor speech-in-noise ability in CI users. Together, these findings may help explain why speech-in-noise outcomes remain highly variable among CI users, even when basic sensory encoding abilities are taken into account.

## Additional Information Competing interests

The authors declare no competing interests.

## Author contributions

Hasan Colak: conceptualisation, study design, data collection, data analysis, data interpretation, manuscript writing

Xiaoxuan Guo: conceptualisation, study design, data interpretation Ester Benzaquén: conceptualisation, study design, data interpretation

Madhusagar Gurusiddappa: conceptualisation, study design, data collection Anirvan Banerjee: conceptualisation, study design

Inyong Choi: conceptualisation, study design, data interpretation William Sedley: conceptualisation, study design, data interpretation

Timothy D. Griffiths: conceptualisation, study design, data interpretation, funding.

## Data Availability Statement

The data that support the findings of this study are available from <DATA will be populated after acceptance>

## Acknowledgments

We thank all participants who volunteered and contributed to this study. We would like thank NIHR for their support during the project. Hasan Colak would also like to thank the Ministry of National Education of the Republic of Türkiye for supporting his PhD studies.

